# Survey of the Australian New Zealand Society for Extracellular Vesicles (ANZSEV) Community

**DOI:** 10.64898/2026.07.06.736235

**Authors:** Liam Hourigan, Lesley Cheng, Kirsty. Danielson

## Abstract

The Australian New Zealand Society for Extracellular Vesicles (ANZSEV) was formally established in 2021 as a regional society for EV researchers. The society holds an annual meeting showcasing world-class research, provides opportunities for local and international networking and collaboration, actively supports a network of ECRs, holds mini-symposiums and provides various opportunities for mentorship and awards for members. As an accompaniment to a recent project to outline the society’s history, a survey of ANZSEV members and its broader network was conducted. Forty individuals responded to the survey with results shining a light on the composition, interests and beliefs of the ANZSEV community.

## Introduction

The Australian and New Zealand Society for Extracellular Vesicles (ANZSEV), is a society of EV researchers based in Australia and New Zealand. It was established in 2021 to formalize the regional EV researcher community, which had previously held annual meetings called the Australasian Extracellular Vesicle Conference. ANZSEV now holds annual meetings in diverse locations across Australia and New Zealand, maintains an active, collaborative community with online presentations and an ECR focused program alongside ther initiatives (Figure 1).

**Figure 1:**
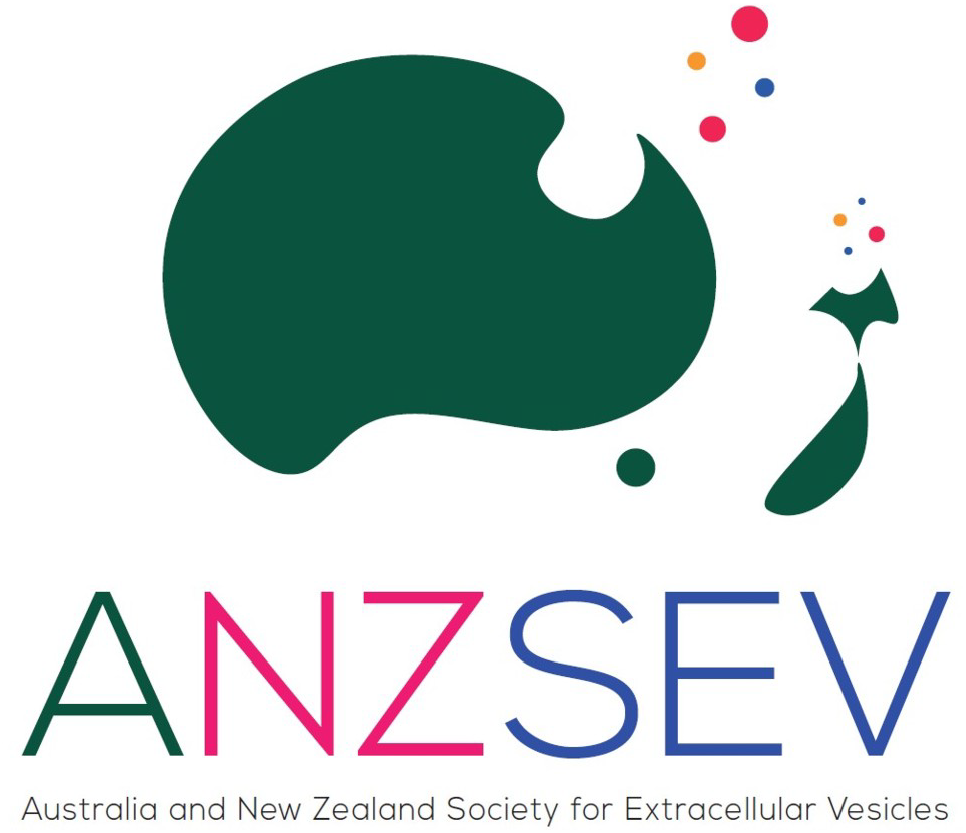
ANZSEV logo. Winning design by Antonia Reale in an open competition held for Australian and New Zealand EV researchers.

In preparation for a profile of the society in an upcoming book, ANZSEV members, under the supervision of President Kirsty Danielson and Treasurer Lesley Cheng, conducted a survey of ANZSEV members and their extended network. The survey aimed to capture insights into the ANZSEV community, on their work, their views of the ANZ EV research community and the broader EV field. This survey, to the author’s knowledge, represents the first formal survey of an EV society.

## Methods

The survey was hosted on the platform Qualtrics and circulated via email and social media to an extended network of over 1300 ANZSEV e-mail list subscribers. The list contained both past and present members, along with other interested individuals, including those from industry. The email requested only those who are members of ANZSEV or based in Australia and New Zealand to respond. The survey was publicly available from 3/5/2026 until 18/5/2026. Full survey questions are available as a supplementary document.

Select ANZSEV survey results are compared against two recent surveys conducted by ISEV. One was conducted by the ISEV Rigor and Standardisation Subcommittee in 2019 and focused on methods for separation and characterization with n=620 total respondents, of which n=320 were complete (Royo et al. 2020). The second ISEV survey was conducted by the Translation, Regulation and Advocacy Committee and focused on their translational potential and industry connections with n=146 total respondents and n=144 complete responses (Gurriaran-Rodriguez et al. n.d.).

## Results and Discussion

### Survey results

ANZSEV had 103 active members at the time of the survey. Forty individuals responded to the survey, of whom 24 were active ANZSEV members (23% of the member base). One ‘other’ respondent stated that they were waiting on funding to initiate membership (Figure 2, Panel A). Raw survey results are available in the associated Open Science Framework repository (Hourigan, 2026). Respondents skewed towards academics (35% of respondents, Lecturer/ Senior Lecturer/Professor), which likely does not reflect the real distribution of ANZSEV members’ career stages (Figure 2, Panel B). ‘Other’ respondents were a research manager and a CEO. This higher engagement from senior academics and researchers was similarly found in surveys conducted by ISEV (Gurriaran-Rodriguez et al. n.d.; Royo et al. 2020). Notably, this ANZSEV-led survey had nearly twice as many ANZ respondents as the 2019 ISEV survey, indicating stronger reach through society-led surveying

**Figure 2:**
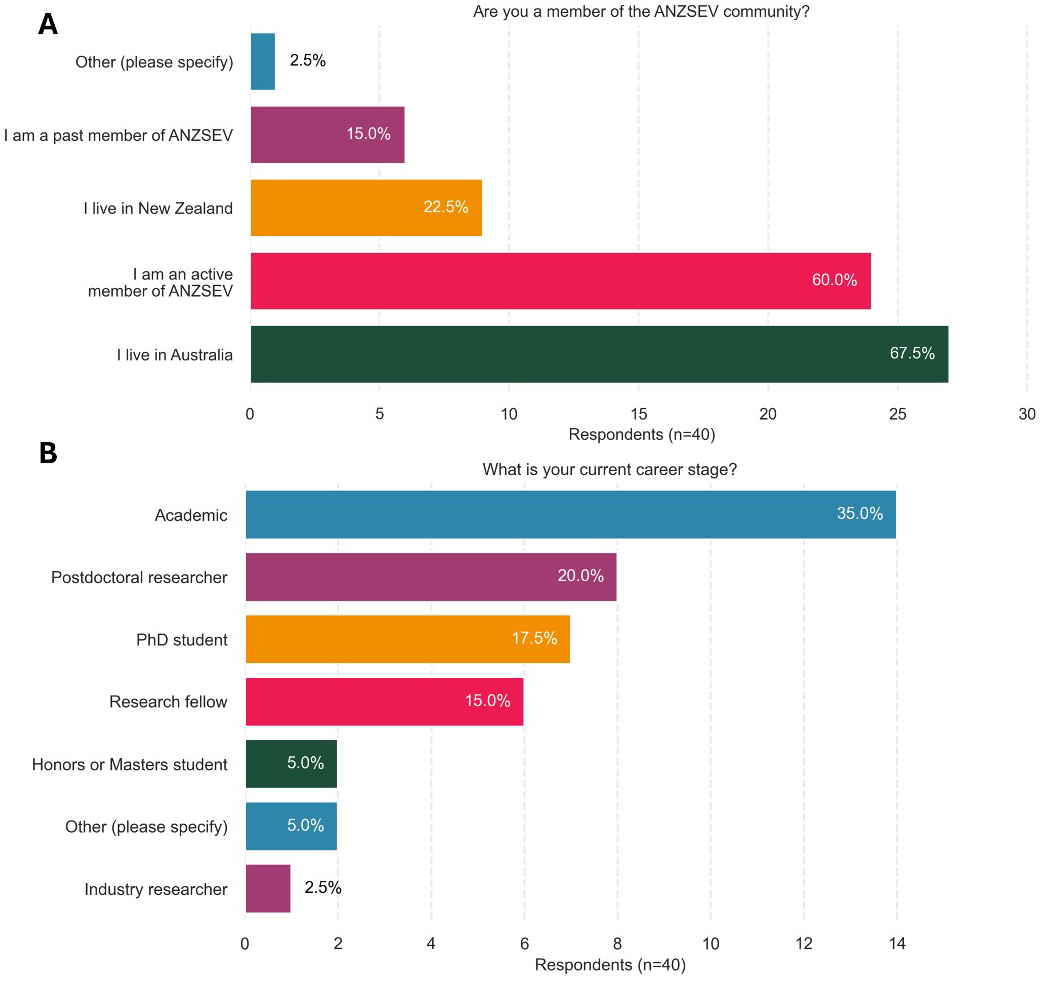
Survey sample characteristics. **(A)** Geographical location, membership status and **(B)** career stage of survey respondents. Academic = (Lecturer/Senior Lecturer/Professor).

Survey Q4 delineated whether researchers considered EV research to be their primary or secondary focus. 23 (58% of total) responded that it was their primary focus, while 16 identified that it was their secondary (40% of total). Twenty-seven respondents provided brief answers detailing their research background before entering the EV field (Q4b). More frequent responses included neurology, stem cell biology, cell biology, microbiology and cancer biology. More diverse responses were also given, including exercise, placental biology, civil engineering, virology and organ-on-a-chip.

Q5 identified whether respondents belonged to an ‘EV research group’. 26 responded ‘yes’ (65%) and 14 responded ‘no’ (35%). Q3 reported the percentage of their research that was EV-focused (Figure 3).

**Figure 3:**
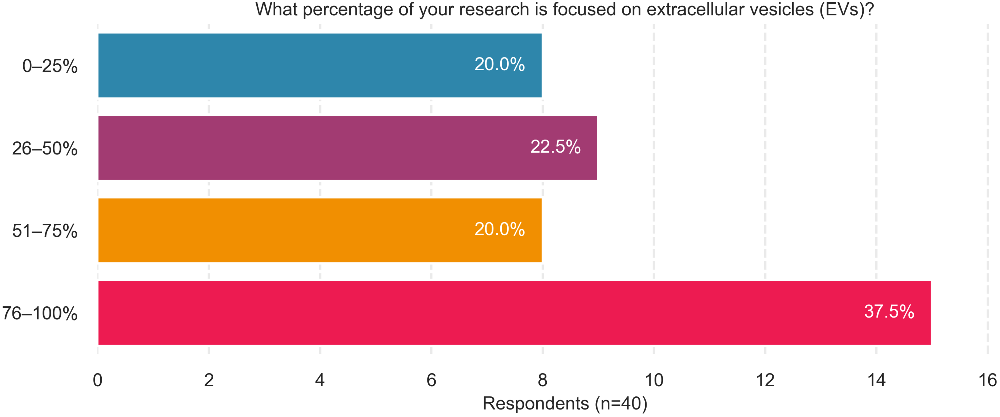
Relative focus on EV research. Individuals gave a singular response for the percentage of their work that was EV-focused. Percentage values within bars reflect totals of n=40 respondents.

Survey Q8 and Q9 asked respondents whether they studied EV function in vivo and/or in vivo. 40% studied function in vitro only, while 42.5% studied function both in vivo and in vitro. No respondents studied in vivo only; one respondent did not answer this question. Comparing these results with the 2019 ISEV methods survey, there appears to be a greater focus on functional assaying among ANZSEV members. In the ISEV survey, 34% investigated functionality both in vitro and in vivo, 34% in vitro only and 3% in vivo. Q6 captured biological sources and models (Figure 4, Panel A), and Q7 identified disease focuses (Figure 4, Panel B).

**Figure 4:**
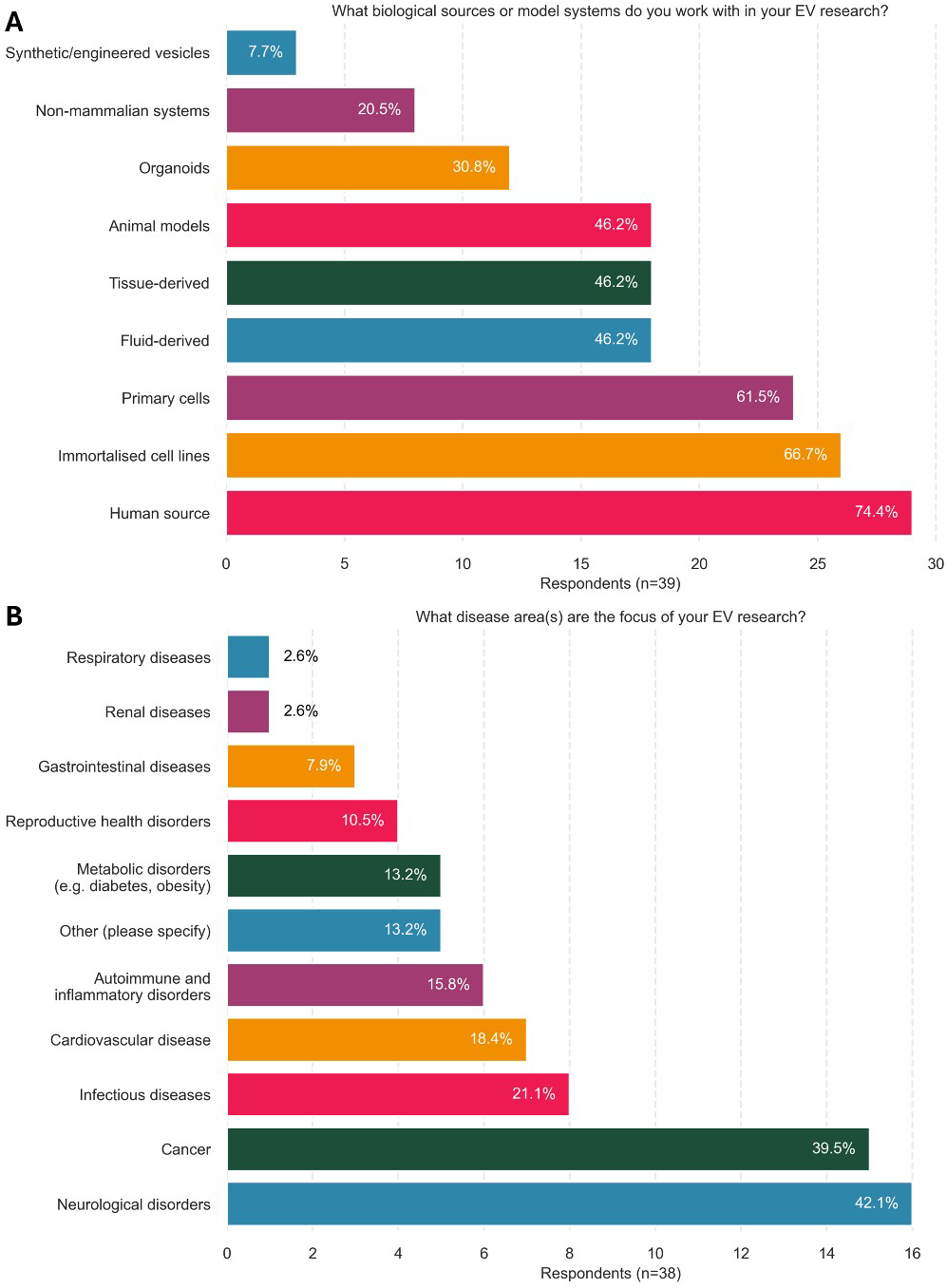
Disease focus and biological sources of ANZSEV researchers. **(A)** Biological sources and models used, and **(B)** Disease research focuses.

Q10 of the survey asked respondents to identify the two categories along a translational pipeline that their research was most focused (Figure 5, panel A). The translational pipeline used was drawn from the Health Research Classifications System (HRCS) used by the United Kingdom Clinical Research Collaboration and adapted for EV research by Hourigan et al. (2025), (Figure 5, panel B). This allows for some comparisons of ANZSEV focus relative to the broader field. However, it should be noted that these results are not directly comparable, as HRCS codings in (Hourigan et al. 2025) reflect publication-level classifications, while the survey captures researcher assessments of their focus. Additionally, as two survey options could be selected, raw percentages can be misleading Nonetheless, some tempered insights can be gleaned by halving the survey percentages to account for double-option selection.

**Figure 5:**
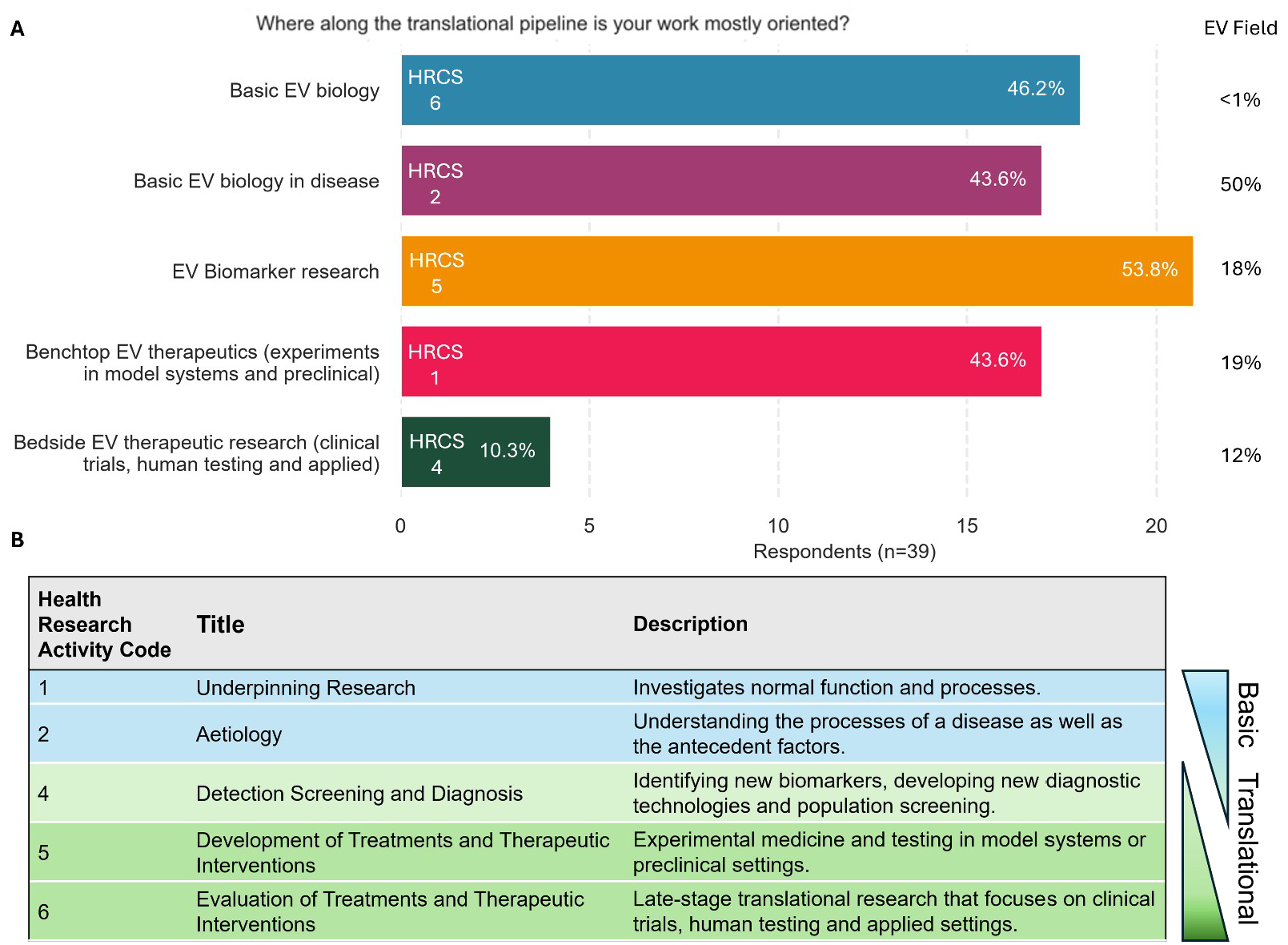
ANZSEV research on the translational pipeline. **(A)** Survey responses placing work focus along the **(B)** Health Research Classifications System (HRCS) translational pipeline. Two options could be selected. Percentages are also given for HRCS classifications of publications in the EV field (Panel A). These reflect the relative amounts of each classification against the whole field (Hourigan et al., 2025).

ANZSEV respondents show a strong focus on biomarker research (halved count 26%) compared to the broader field (12%). Additionally, ‘bedside’ research (halved count 5%) was fivefold more than the broader field (<1%). This may reflect the senior academic leaning survey respondents and strong ties to industry and contact with regulatory agencies. A 10% involvement in EV-related clinical trials was also found in the 2024 TRA survey which sampled a similar population of industry-tied academics.

### Industry connections

ANZSEV survey respondents indicated strong translational leanings with Q11 asking whether individuals had made any contact with regulatory agencies about clinical EV applications. Nine (22.5%) responded ‘yes’, with Q12 asking whether individuals had connections with industry. 14 (35%) responded ‘yes’ with these connections further detailed in Figure 6.

**Figure 6:**
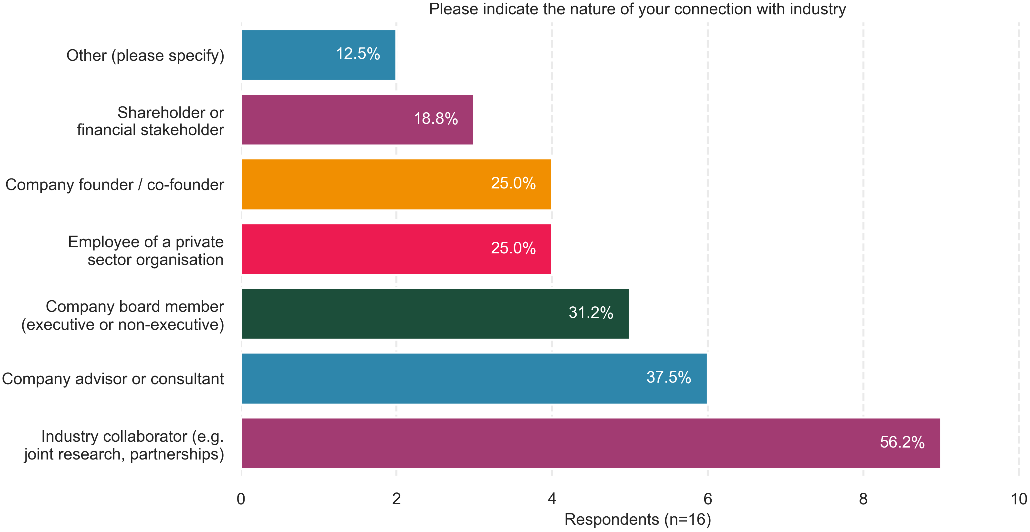
Industry connections. Industry connections of survey respondents. Note that percentages reflect respondents to this question (n=16) not the overall survey (n=40). Respondents could provide more than one answer.

Some comparisons can be made against the 2024 ISEV Translation, Regulation and Advocacy Committee (ISEV-TRA) survey (Gurriaran-Rodriguez et al. n.d.). Their survey included n=144 respondents and was also skewed towards more senior researchers and academics, with both surveys indicating a high degree of industry engagement among surveyed senior EV researchers. The ISEV survey identified 12 respondents (8%) serving on an advisory board, while the ANZSEV survey identified 6 respondents (15% of n=40) serving on an advisory board or as consultants.

### Methods used by ANZSEV members

ANZSEV survey responses for methods used are largely comparable to the ISEV 2019 methods survey, as questions included mostly the same methods. However, statistical comparisons could not be made due to a lack of access to ISEV data. Figure 7 provides a comparison of isolation techniques. The most notable differences between the ANZSEV and ISEV surveys are an emphasis on size exclusion chromatography (SEC) and tangential flow filtration (TFF) in ANZSEV respondents. This difference may partly reflect favoured methods within ANZSEV and/or the fact that SEC and TFF have become more widely used over time. ‘Other’ respondent uses FACS based sorting and differential centrifugation.

**Figure 7:**
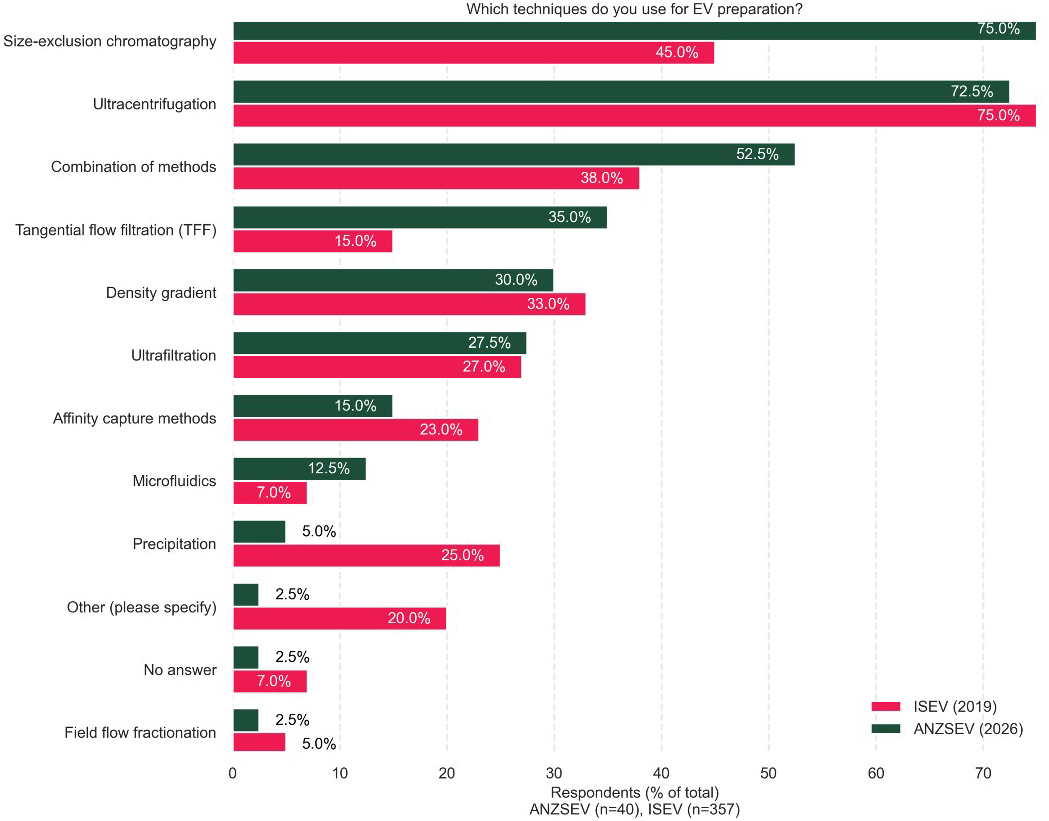
Isolation methods. Isolation techniques of ANZSEV and ISEV survey respondents Results from 2019 ISEV survey have a ±2-3% margin of error as numbers were visually extracted from figures due to unclear reporting.

Figure 8, panel A, covers characterisation methods and includes some differences to the ISEV survey in the methods represented. The ISEV survey includes atomic force microscopy and surface plasmon resonance, while the ANZSEV survey includes RNA concentration and tunable resistive pulse sensing. Fewer substantial differences were observed, however, there does appear to be a greater usage of electron microscopy, both cryo and non-cryo, among ANZSEV respondents. Figure 8, panel B compares the number of methods used against the distribution of the same chart in the ISEV survey. The ISEV chart appears more normally distributed (likely due to their higher number of respondents), centred around a 4-6 method peak. ANZSEV respondents mostly used 4-5 methods, with another small peak at eight methods.

**Figure 8:**
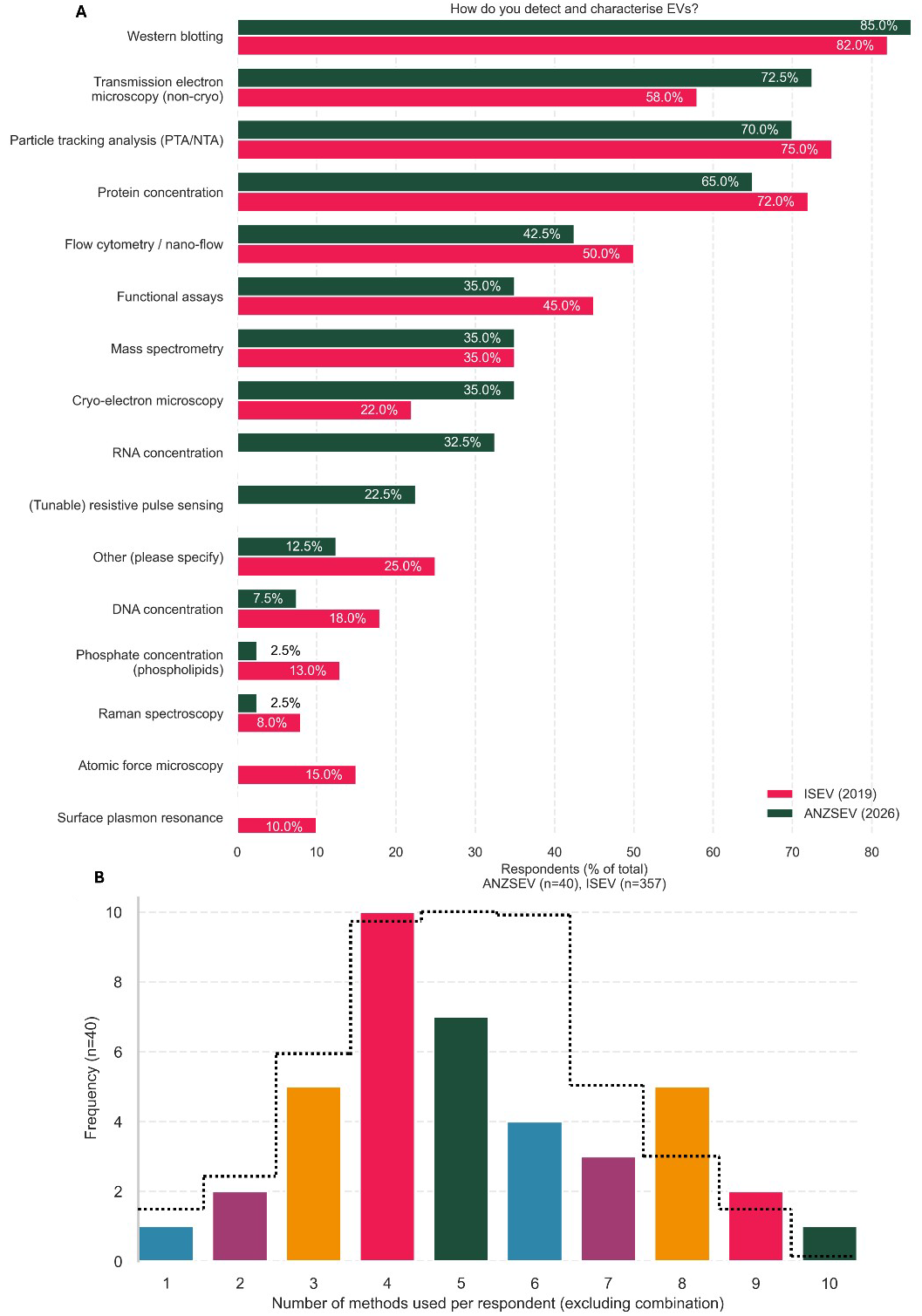
Characterization methods. **(A)** Isolation techniques of ANZSEV (n=40) and ISEV (n=357) survey respondents and **(B)** number of methods used per ANZSEV and ISEV respondent. Results from 2019 ISEV survey (black lines) have a small margin of error, as relative bar sizes were visually extracted from Figure 6 due to unclear reporting . ISEV results are not scaled to the y-axis, they are a distributional representation.

### Views on standardization and replication

The close ties between the ANZSEV research community and ISEV during its formative years have seemingly made a strong impression on ANZSEV, with 72% of respondents agreeing that ANZSEV places a high value on rigour and standardisation, while 21% were unsure. Respondents gave short answers on what ‘strong rigour and standardisation meant for them’. Most respondents emphasise reproducibility and transparency in reporting findings. One gave a detailed response “Academic standards for rigor and standardisation are very different from the industry standard of rigor and standardisation. I still find that publications on EVs don’t have enough detail to replicate their findings. Also, small changes can affect EV production and EV function, and the standards for academia don’t seem to be the same as they are for industry, for things like cell culture, acceptance of reagents, lot numbers, expiry dates, etc. I am unsure about how this would affect research outcomes.”. Other respondents echoed this sentiment: reproducibility in basic research was both a necessity and a current handicap to further translation. Two respondents mentioned ISEV guidelines with one showing support and another expressing disagreement.

Short answers were also given explaining the most important challenge the field faces, with 30% of respondents mentioning standardisation, while 20% identified reproducibility. Issues were flagged in basic EV science, including concerns about purity and more specifically that some previously reported EV functions may be attributable to co-isolates. Many responses centred on ‘success’ around the ultimate goal of clinical translation, identifying concerns such as rigour, standardisation, and reproducibility, as well as gaps in basic research that need to be addressed to achieve this. Some mentioned more industry-associated terms, such as manufacturing methods and scalability.

The ANZSEV survey asked respondents about their experiences in replicating findings (Figure 9). To our knowledge, this is the first formal survey of replication experiences within the EV field. Thus, there are no within-field points of comparison. However, a rough comparison can be made against a survey on reproducibility perspectives of researchers in biomedicine (Cobey et al. 2024). Among their n=1630 respondents, 47% reported failing to replicate published findings of another group. This closely aligns with the 43% of total ANZSEV survey respondents (n=40) who answered a similar question.

**Figure 9:**
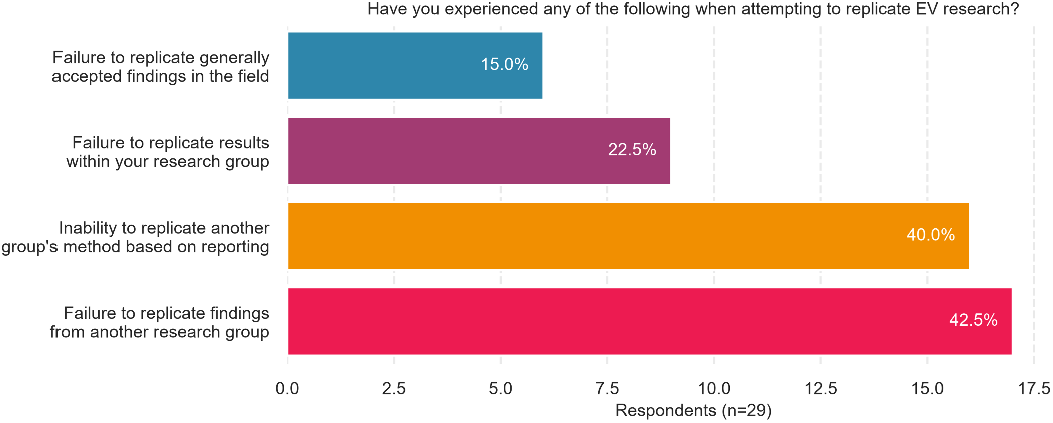
Replication in the EV field. Reported challenges in replication. Percentages calculated based on question respondents (i.e. n=29).

### ANZSEV respondent perspectives on ANZ EV research

Q16 asked survey respondents, “What do you see as the key strengths or central focuses of EV research in Australia/New Zealand?” Many responses cited a strong collaborative culture and a community of practice which shares information and provides support. Two extensive responses are reflective of the community’s placement on the translational pipeline. “Diversity, support, collaborative approach for infrastructure, access to clinical cohorts, data informatics and machine modelling labs and resources, extensive background of researchers across regional and international levels in EV research, expertise, journals, international guidelines, task forces and other major activities of EV field at international level” and “Strong translational focus, particularly using EVs as liquid biopsy tools for clinically relevant questions. Growing strength in bioengineering and EV-based therapeutics. A highly collaborative research network”

Q18 asked survey respondents what they would “like to see more of from ANZSEV in the future” with 24 respondents giving at least one suggestion. A synthesis of suggestions is provided:

- Database of resources, including experts for support/collaboration and equipment.
- Opportunities to form interdisciplinary connections and develop skills through workshops and networking events with experts of diverse backgrounds.
- More online symposia. International speakers and/or ISEV collaboration.
- Smaller regional gatherings/events with casual presentations and ECR opportunities
- ECR/student mentoring and coaching opportunities
- Public outreach and science communication of EV research
- Hands on workshops for clinical translation, regulatory pathways, GMP manufacturing, and entrepreneurship. Providing opportunities for industry and regulatory connections.
- Advocating for EV research and application with governments and regulatory agencies.
- ANZSEV led taskforces and working groups
- Technology updates and opportunities to trial new technologies
- Workshops/opportunities to develop large collaborative projects and grants.

### Perspectives on EV field maturity

The survey asked respondents to rate how ‘mature’ they believed the EV field to be, with a follow up short answer explaining their response (Figure 10). The rating was a 1-10 scale with 1 = not mature to 10 = very mature with no other context given. Respondents mostly indicate that the field is at a midway point, based on their own conceptions of maturity. What follows is a synthesis of these responses.

**Figure 10:**
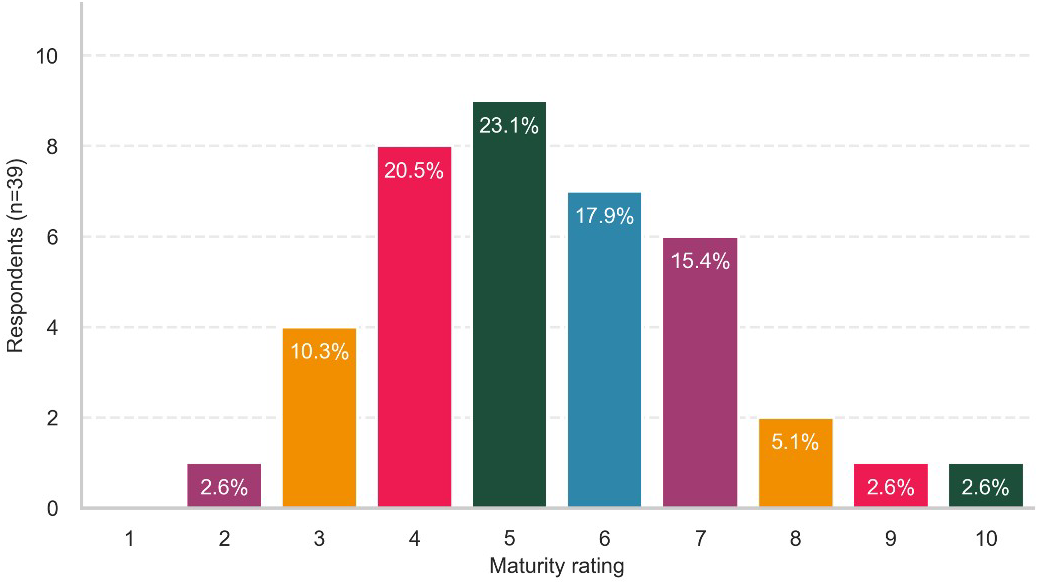
Maturity. Respondents answered “how mature is the EV field” on a 10-point scale.

The previous question on challenges facing the field highlighted that many researchers centre clinical applications as the endpoint of research efforts. This is reflected in ‘maturity’ responses, with this ‘midpoint’ of maturity based on a growing understanding of basic biology and a lack of later translational success or ‘poster stories’. The field has passed through the early promise and excitement of EVs, and has reached a “plateau of novelty”. The explosion of EV research over the last decade is now being matched by newer, more precise technologies. These allow the field to begin answering more long-standing questions, while also uncovering issues with past findings and reproducibility. There is also heterogeneity in knowledge about subtypes of basic EVs. Small mammalian-derived EVs are the most studied and understood, while large EVs and those from more diverse organisms are less so. Additionally, it is noted that the EV function is mostly understood in vitro, while a more complex systems-level understanding of EVs in physiology remains patchy. In turn, it seems recognised that the field has grappled with classification through multiple MISEV guidelines, and progress is being made.

On the translational side, the lack of successful stories and the irreproducibility of ‘dodgy’ translational claims were highlighted. The Good Manufacturing Practice (GMP) and regulatory pathways were flagged as immature. One respondent stated, “LNPs (lipid nanoparticles) have completely overtaken EVs in the last 10 years in respect of the size of the field and number of applications. Also, the largest current uptake, cosmetics, is not really science-based”.

## Conclusion

Our survey captured the views of almost a quarter of active ANZSEV members as well as those of the broader ANZSEV network. Our sampling was skewed towards senior academics, many of whom had connections with industry. 58% of respondents had at least a minimum of 50% focus on EV research. This is an interesting insight, as many researchers working with EVs have transitioned from other research backgrounds and may continue to operate in this area. To date, this relative focus has not been quantified. Without reference to other areas, it is unclear whether ANZ EV researchers are any more or less solely focused on EVs than those of other countries.

We compared some of our findings against two other ISEV surveys, noting some differences in the methods used. ANZ researchers present SEC and TFF as favoured isolation methods, alongside electron microscopy, for characterisation. We also found that our ANZ respondents had greater involvement in clinical trials and certain industry positions than those in the ISEV TLA survey.

Our survey also highlighted a gap in formal knowledge around perceptions of reproducibility. Reproducibility was frequently highlighted as a major challenge facing the field, yet our survey was the first to ask respondents directly about their experiences with replication. Having 15% of total respondents identify issues with replication of generally accepted findings in the field is concerning. In survey on reproducibility in biomedicine by (Cobey et al. 2024) they identified that 27% of their respondents believed there was a significant ‘crisis’ in reproducibility, while 45% believed there was a ‘slight’ crisis. It would be valuable to determine, in a more widespread survey, whether EV researchers believe the field is undergoing a crisis similar to that of other related fields.

We captured valuable insights into ANZ EV researchers’ beliefs around their field’s maturity as they conceived it. Many detailed responses were given explaining assessments of maturity, indicating that this is an interesting topic or point of discussion for EV researchers. Most placed the EV field at a midpoint, one that was based on a growing understanding of basic EV biology yet falling short of substantial translational success.

## Supporting information

Survey Questions

## Acknowledgements

David Greening (past president of ANZSEV) was supportive of this project and William Phillips tested an early draft of the survey and provided feedback.

## Author contributions

CRediT contributions: Liam Hourigan (Conceptualization, Investigation, Formal analysis, Methodology, Writing – original draft); Lesley Cheng (Methodology, Writing – review and editing) and Kirsty Danielson (Methodology, Supervision, Writing - review and editing).

