## Supplementary material for "Survey of the Australian New Zealand Society for Extracellular Vesicles (ANZSEV) Community": Survey Questions

### Survey of ANZSEV Community

Start of Block: Section 1 - Background and Research Profile - Apr 20, 2026

Q1 Are you a member of the ANZSEV community?

- ☐ I live in Australia (1)
- ☐ I live in New Zealand (2)
- ☐ I am an active member of ANZSEV (3)
- ☐ I am a past member of ANZSEV (4)
- ☐ Other (please specify) (5) __________________________________________________

Q2 What is your current career stage?

- ○ Honors or Masters student (1)
- ○ PhD student (2)
- ○ Postdoctoral researcher (3)
- ○ Research fellow (4)
- ○ Academic (Lecturer/Senior Lecturer/Professor) (5)
- ○ Clinician-researcher (6)
- ○ Industry researcher (7)
- ○ Other (please specify) (8) __________________________________________________

Q3 What percentage of your research is focused on extracellular vesicles (EVs)?

- ◯ 0–25%  (1)
- ◯ 26–50%  (2)
- ◯ 51–75%  (3)
- ◯ 76–100%  (4)

---

Q4 How would you describe your involvement in EV research?

- ◯ I am an EV researcher (primary focus)  (1)
- ◯ I am a researcher working with EVs (secondary focus)  (2)

---

Q4b Briefly, what was your research background prior to working with EVs? (Leave blank if not applicable)

________________________________________________________________

---

Q5 Would you describe yourself as being based in an 'EV research group'?

- ◯ Yes  (1)
- ◯ No  (2)

**End of Block: Section 1 - Background and Research Profile - Apr 20, 2026**

---

**Start of Block: Section 2 - Research Characteristics - Apr 20, 2026**

Q6 What biological sources or model systems do you work with in your EV research? (Select all that apply)

- ☐ Human source (1)
- ☐ Animal models (e.g. mouse, rat, zebrafish) (2)
- ☐ Non-mammalian systems (e.g. plants, bacteria, parasites, fungi) (3)
- ☐ Tissue-derived (4)
- ☐ Fluid-derived (5)
- ☐ Cell culture (immortalised cell lines) (6)
- ☐ Primary cells (7)
- ☐ Organoids (8)
- ☐ Synthetic/engineered vesicles (9)
- ☐ Other (please specify) (10) ________________________________________________

Q7 What disease area(s) are the focus of your EV research? (Select all that apply)

- ☐ Cancer (1)
- ☐ Cardiovascular disease (2)
- ☐ Neurological disorders (3)
- ☐ Infectious diseases (4)
- ☐ Autoimmune and inflammatory disorders (5)
- ☐ Metabolic disorders (e.g. diabetes, obesity) (6)
- ☐ Respiratory diseases (7)
- ☐ Renal diseases (8)
- ☐ Gastrointestinal diseases (9)
- ☐ Reproductive health disorders (10)
- ☐ Rare/genetic diseases (11)
- ☐ Other (please specify) (12) ________________________________________________

---

Q8 Do you study EV function in vitro?

- ○ Yes (1)
- ○ No (2)

---

Q9 Do you study EV function in vivo?

○ Yes (1)

○ No (2)

Q10 **Figure.** Hourigan, L., Phillips, W., Kenari, A. N., Pavani, K. C., Chen, C., Hendrix, A., Cheng, L., & Hill, A. F. (2025). Mapping growth and trajectory in the field of extracellular vesicles: A scientometric analysis. *Extracellular Vesicle*, *5*, 100062. https://doi.org/https://doi.org/10.1016/j.vesic.2024.100062 Where along the translational pipeline is your work mostly oriented? (Select up to 2)

▢ Basic EV biology (Underpinning research) (1)

▢ Basic EV biology in disease (Aetiology) (2)

▢ EV Biomarker research (Detection, screening and diagnosis) (3)

▢ 'Benchtop' EV therapeutic research e.g. experiments in model systems and preclinical settings (Development of treatments and therapeutic interventions) (4)

▢ 'Bedside' EV therapeutic research e.g. clinical trials, human testing and applied settings (Evaluation of treatments and therapeutic interventions) (5)

Q11 Do you have contact with regulatory agencies regarding clinical application of EVs?

○ Yes (1)

○ No (2)

End of Block: Section 2 - Research Characteristics - Apr 20, 2026

Start of Block: Section 3 - Industry Connections - Apr 20, 2026

Q12 Do you have any connection with industry (e.g. company board member)?

- Yes (1)
- No (2)

Q12a Please indicate the nature of your connection with industry: (Select all that apply) *Note: Complete this question only if you answered Yes above.*

- Company founder / co-founder (1)
- Company board member (executive or non-executive) (2)
- Company advisor or consultant (3)
- Employee of a private sector organisation (4)
- Industry collaborator (e.g. joint research, partnerships) (5)
- Shareholder or financial stakeholder (6)
- Other (please specify) (7) ________________________________________________

Q12b Is this connection related to your EV research activities? *Note: Complete this question only if you answered Yes to Q12 above.*

- Yes, directly related (1)
- Yes, indirectly related (2)
- No (3)

**End of Block: Section 3 - Industry Connections - Apr 20, 2026**

**Start of Block: Section 4 - Methods and Techniques - Apr 20, 2026**

Q13 Which techniques do you use for EV preparation? (Select all that apply)

- ▢ Affinity capture methods (1)
- ▢ Density gradient (2)
- ▢ Field flow fractionation (3)
- ▢ Microfluidics (4)
- ▢ Precipitation (5)
- ▢ Size-exclusion chromatography (6)
- ▢ Ultracentrifugation (7)
- ▢ Ultrafiltration (8)
- ▢ Tangential flow filtration (TFF) (9)
- ▢ Combination of methods (10)
- ▢ Other (please specify) (11) ________________________________________________

Q14 How do you detect and characterise EVs? (Select all that apply)

- ▢ Atomic force microscopy (1)
- ▢ Cryo-electron microscopy (2)
- ▢ DNA concentration (3)
- ▢ Flow cytometry / nano-flow (4)
- ▢ Functional assays (5)
- ▢ Mass spectrometry (6)
- ▢ Particle tracking analysis (PTA/NTA) (7)
- ▢ (Tunable) resistive pulse sensing (8)
- ▢ Phosphate concentration (phospholipids) (9)
- ▢ Protein concentration (10)
- ▢ Raman spectroscopy (11)
- ▢ RNA concentration (12)
- ▢ Surface plasmon resonance (13)
- ▢ Transmission electron microscopy (non-cryo) (14)
- ▢ Western blotting (15)
- ▢ Other (please specify) (16) ________________________________________________

Q15 Have you experienced any of the following when attempting to replicate EV research? (Select all that apply)

▢ Failure to replicate results within your research group (1)

▢ Failure to replicate findings from another research group (2)

▢ Failure to replicate generally accepted findings in the field (3)

▢ Inability to replicate another group's method based on reporting (4)

**End of Block: Section 4 - Methods and Techniques - Apr 20, 2026**

---

**Start of Block: Section 5 - EV Research in Australia and New Zealand - Apr 20, 2026**

Q16 What do you see as the key strengths or central focuses of EV research in Australia/New Zealand?

________________________________________________________________

________________________________________________________________

________________________________________________________________

________________________________________________________________

________________________________________________________________

---

Q17 Do you believe the Australia/New Zealand EV research community places a high value on rigor and standardisation?

○ Yes (1)

○ No (2)

○ Unsure (3)

---

Q17b What does strong rigor and standardisation mean to you? (Optional)

________________________________________________________________

________________________________________________________________

________________________________________________________________

________________________________________________________________

________________________________________________________________

Q18 What would you like to see more of from ANZSEV in the future? Please list as many ideas as you like (e.g. initiatives, events, resources, collaborations).

- ○ Idea 1 (1) ________________________________________________
- ○ Idea 2 (2) ________________________________________________
- ○ Idea 3 (3) ________________________________________________
- ○ Idea 4 (4) ________________________________________________
- ○ Idea 5 (5) ________________________________________________

**End of Block: Section 5 - EV Research in Australia and New Zealand - Apr 20, 2026**

**Start of Block: Section 6 - Your Views on the EV Field - Apr 20, 2026**

Q19 What is the most important challenge facing the EV field globally?

________________________________________________________________

________________________________________________________________

________________________________________________________________

________________________________________________________________

________________________________________________________________

Q20 How "mature" is the EV field?

| | 1 – Newly founded (1) | 2 (2) | 3 (3) | 4 (4) | 5 (5) | 6 (6) | 7 (7) | 8 (8) | 9 (9) | 10 – Very mature (10) |
|---|---|---|---|---|---|---|---|---|---|---|
| Maturity rating (1) | ○ | ○ | ○ | ○ | ○ | ○ | ○ | ○ | ○ | ○ |

Q20b Please explain your maturity rating above. (Optional)

________________________________________________________________

________________________________________________________________

________________________________________________________________

________________________________________________________________

________________________________________________________________

**End of Block: Section 6 - Your Views on the EV Field - Apr 20, 2026**
